# Hepatobiliary organoids derived from leporids support the replication of hepatotropic lagoviruses

**DOI:** 10.1101/2022.04.07.487566

**Authors:** Egi Kardia, Omid Fakhri, Megan Pavy, Hugh Mason, Nina Huang, Elena Smertina, Mary K. Estes, Tanja Strive, Michael Frese, Robyn N. Hall

## Abstract

Lagoviruses, family *Caliciviridae*, are some of the most virulent vertebrate viruses known. They infect leporids, i.e., rabbits (*Oryctolagus* spp.), hares (*Lepus* spp.) and cottontails (*Sylvilagus* spp.) and rapidly kill over 95% of susceptible animals. Pathogenic lagoviruses are hepatotropic and induce a fulminant hepatitis that typically leads to disseminated intravascular coagulation within 24–72 hours of infection. However, the pathophysiological mechanisms governing this extreme phenotype and other aspects of the fundamental biology of these viruses are poorly understood due to a lack of cell culture systems. Here, we report on a robust and reliable *ex vivo* model for the cultivation of hepatotropic lagoviruses. We show that three rabbit haemorrhagic disease virus (RHDV) variants, RHDV1, RHDV2 and RHDVa-K5, replicate in monolayer cultures derived from rabbit hepatobiliary organoids. Viral replication was demonstrated by a (*i*) increase in viral RNA levels of greater than one log_10_ over a 23-hour period, (*ii*) detection of viral structural and non-structural proteins, and (*iii*) detection of double-stranded RNA viral replication intermediates. Furthermore, we generated hepatobiliary organoids from a feral cat (*Felis domesticus*), a wild mouse (*Mus musculus*) and a European brown hare (*Lepus europaeus*) and showed that monolayer cultures derived from the cat and mouse organoids were not permissive for lagovirus infection, while those derived from the hare organoids only supported replication of RHDV2, recapitulating the species tropism that has been observed with these viruses. Our organoid culture system will facilitate future studies into the molecular biology of lagoviruses, which will have considerable import for the conservation of endangered leporid species in Europe and North America and the biocontrol of overabundant rabbit populations in Australia and New Zealand.

## Introduction

Lagoviruses are caliciviruses that infect leporids, i.e., rabbits (*Oryctolagus* spp.), hares (*Lepus* spp.) and cottontails (*Sylvilagus* spp.), and are some of the most virulent vertebrate-infecting viruses known, with some genotypes leading to death in over 95% of susceptible animals [1, 2]. All known pathogenic lagoviruses are hepatotropic and cause massive hepatic necrosis, disseminated intravascular coagulation, multi-organ failure and death within 24–72 hours of infection [2, 3]. Several enterotropic lagoviruses have also been identified but have not been associated with clinical disease. While some variants such as RHDV1 and RHDVa (genotypes GI.1 and GI.1a, respectively [4]) only infect European rabbits (*Oryctolagus cuniculus*), RHDV2 (genotype GI.2) has been shown to also infect several hare, jackrabbit and cottontail species of the genera *Lepus* and *Sylvilagus* [5–12]. This host-range expansion beyond that recognised for the GI.1 lagoviruses, coupled with the panzootic spread of RHDV2 over the last decade, has led to concerns that this virus may infect other domestic species or endangered native wildlife. Unfortunately, the mechanisms that govern host and tissue tropism, virulence and other fundamental biological properties of lagoviruses are currently poorly understood due to lack of a robust cell culture system for these viruses.

Like many other caliciviruses, any *ex vivo* cultivation of lagoviruses has so far proven challenging. While RHDV1 can be cultivated in primary rabbit hepatocytes, these cultures are laborious to establish and can only be maintained for two to three weeks [13]. One research group has reported the establishment of several reverse genetics systems for RHDV1; however, there are no reports of these systems being successfully replicated in other laboratories [14–17]. Organoid and organoid-derived culture systems offer a novel avenue to cultivate lagoviruses *ex vivo*, following on from the discovery that human enteric organoids support the replication of human caliciviruses (noroviruses) [18].

Organoids are three-dimensional, non-transformed, self-organised, multicellular tissue constructs that mimic the cellular architecture and composition of the corresponding organ. Since pathogenic lagoviruses are hepatotropic, a biologically relevant culture system should be representative of the hepatic parenchyma and hepatobiliary tract (Figure 1). Briefly, the liver is comprised of two major cell populations: hepatocytes or parenchymal cells constitute 70–80% of the liver mass and 60% of cell numbers, while cholangiocytes (biliary epithelial cells) represent 3–5% of liver cells [19, 20]. The remaining cell populations comprise hepatic stellate cells, the sinusoidal endothelium, Kupffer cells, specialised hepatic natural killer (NK) cells and itinerant leukocytes [19, 20]. Hepatic organoids from several species (humans, mice, dogs, pigs, cats, chickens and trout [21–29]) have been generated previously, either from primary biopsy specimens containing hepatic progenitor cells or via the induction of pluripotent stem cells. Here we report the successful cultivation of hepatotropic lagoviruses in non-transformed monolayer cultures derived from rabbit hepatobiliary organoids, while virus replication was not supported in cat or mouse hepatobiliary organoid-derived monolayer cultures.

**Figure 1:**
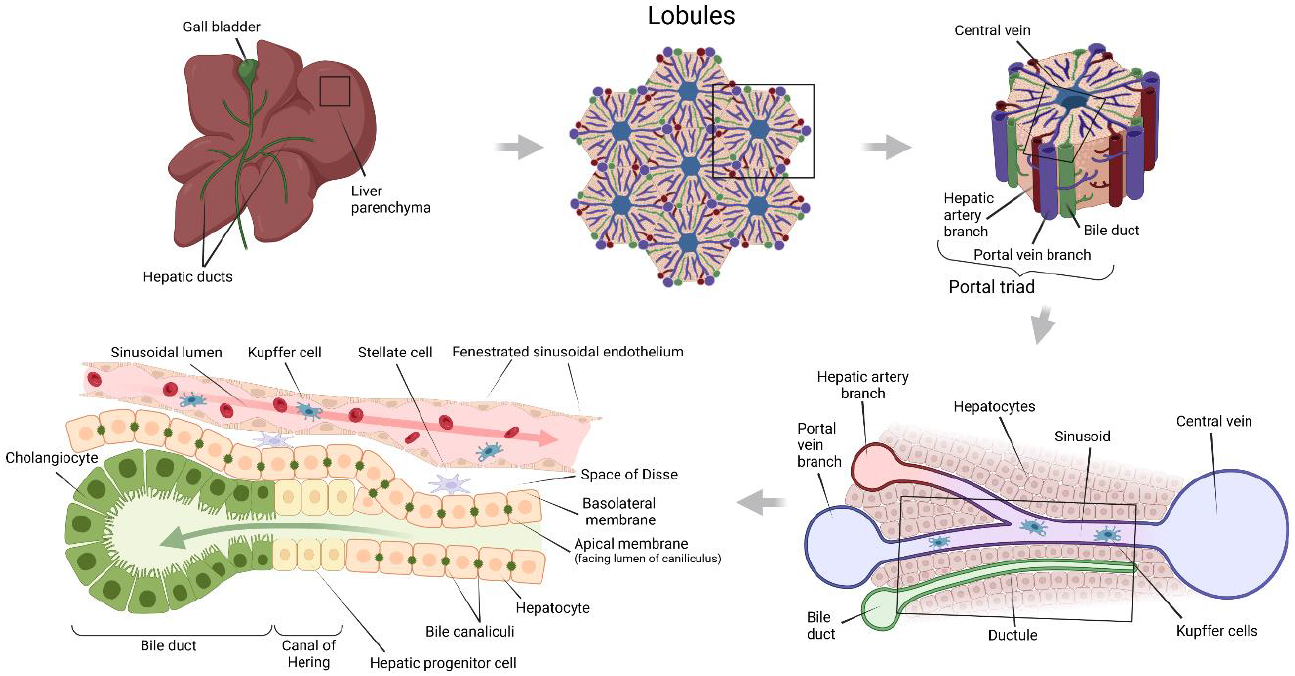
Schematic view of the hepatic functional unit and hepatobiliary progenitor cells. The liver is broadly comprised of the parenchyma (liver lobes) and the biliary tree (gall bladder and associated hepatic ducts). At a microscopic level, the liver parenchyma is arranged into lobules or acini. Lobules comprise a hexagonal arrangement of hepatocytes centred on the terminal hepatic venule or central vein and outlined by fibrovascular septae extending between portal triads (portal tracts), comprising a portal venule, hepatic arteriole and bile duct surrounded by connective tissue. Within the lobule, single-cell-thick anastomosing plates of hepatocytes are separated by hepatic sinusoids, which supply blood to the parenchyma. At their basolateral surface, hepatocytes interface with a perivascular space (space of Disse) containing stellate cells, reticulin fibres and nerves; this space also interfaces with the specialised fenestrated sinusoidal endothelium that facilitates exchange of metabolites with the blood. Hepatocytes secrete bile acids into the bile canaliculus via their apical surface. Bile moves through the canaliculi into the canal of Hering and then into bile ductules (cholangioles) and the interlobular bile ducts. Hepatic progenitor cells reside in the canal of Hering; these progenitors are bipotential, i.e., they can mature into either cholangiocytes or hepatocytes. As such, they express markers of both cell types (e.g., cytokeratin-19 (KRT-19) and albumin (ALB), respectively). The figure was created with BioRender.com.

## Methods

### Isolation and maintenance of 3D rabbit hepatobiliary organoids

Hepatic progenitor cells were isolated from one healthy 7-week-old male New Zealand white rabbit (*Oryctolagus cuniculus*; CSIRO Wildlife and Large Animal - Animal Ethics Committee [CWLA-AEC] permit number 2020-01). The animal was humanely killed by intravenous injection of Lethabarb (Virbac) after intramuscular sedation with a combination of 30 mg/kg ketamine (Mavlab) and 5 mg/kg xylazine (Troy Laboratories). The isolation of hepatic progenitor cells was performed as described by Broutier *et al*. [28, 30], with modifications. Briefly, a section of liver was collected from the central hilar region and placed into basal medium (Supplementary Table 1). Liver was minced and rinsed in wash medium (Supplementary Table 1). Tissue pieces were resuspended in digestion medium (Supplementary Table 1) and incubated at 37°C for 60 minutes with gentle shaking. The remaining liver fragments were transferred into a Petri dish and a scalpel blade was used to scrape away the remaining parenchyma and expose the bile duct structure. The bile duct fragments were further digested in 0.25% trypsin/EDTA (Thermo Fisher Scientific) for 10 minutes at 37°C and passed through a 70-μm cell strainer. The digestion was stopped by adding basal medium containing 20% fetal bovine serum (Thermo Fisher Scientific) and the cell suspension was pelleted by centrifugation at 200 × g for 5 minutes at 4°C and washed. Red blood cells were removed using Hybri-Max red blood cell lysis buffer (Sigma-Aldrich) and the cells were pelleted again by centrifugation at 200 × g for 5 minutes at 4°C. The epithelial cells were then washed twice with phosphate buffered saline (PBS), pelleted, and resuspended in thawed Geltrex lactose dehydrogenase elevating virus free (LDEV-free) reduced growth factor basement membrane matrix (Thermo Fisher Scientific). Three 10-μl drops of the matrix-cell suspension were pipetted into wells of a Nunclon Delta 24-well plate (Thermo Fisher Scientific) and were solidified for 15 minutes at 37°C before adding 400 μl per well of culture medium (Supplementary Table 1). The cultures were incubated at 37°C in 5% CO_2_ and monitored daily to assess the formation of organoids. The medium was changed every 4 days and organoids were sub-cultured when the matrix dome became crowded. Passaging and cryopreservation of the culture was performed as described previously [31].

### Hepatobiliary organoid-derived monolayer culture

Liver organoids were transformed into 2D monolayer cultures in either flat-bottom tissue culture-treated 96-well plates (Corning) or in 8-well Nunc Lab-TekII chamber slides (Thermo Fisher Scientific), for RNA isolation or microscopy, respectively. Prior to use, both plates and chamber slides were coated with 1:50 Geltrex-LDEV (Thermo Fisher Scientific) in PBS for at least 2 hours at 37°C. Proliferating organoids were grown for 4–7 days and dissociated using TrypLE Express Enzyme (Thermo Fisher Scientific) for 10–20 minutes in a 37°C water bath. Basal medium containing 10% FBS was added, and cells were pelleted and resuspended in culture medium. Approximately 7 × 10^4^ cells in 100 μl or 1.4 × 10^5^ cells in 200 μl were plated per well in a 96-well plate or an 8-well chamber slide, respectively.

### Hare-, cat- and mouse-derived hepatobiliary organoid monolayer cultures

European brown hare (*Lepus europaeus*) and cat (*Felis catus*) liver samples were collected opportunistically during routine feral animal control operations in New South Wales, Australia (CWLA-AEC permit number 2019-28). A mouse (*Mus musculus*) liver sample was collected opportunistically through an existing rodent sampling project (CWLA-AEC permit number 2020-24). All animals were presumed to be healthy. Liver samples were processed in the field as described above and transported to the laboratory in cold basal medium (Supplementary Table 1). Hepatic progenitor cells were isolated and propagated as for the rabbit hepatobiliary organoids, described above.

### Viruses

Three lagoviruses were used in this study: RHDV1 (genotype GI.1cP-GI.1c), RHDVa-K5 (genotype GI.1aP-GI.1a) and RHDV2 (genotype GI.1bP-GI.2). Genome sequences are available under GenBank accession numbers KT344770, MF598301 and MW467791, respectively. Briefly, freeze-dried virus stocks were prepared by the Elizabeth Macarthur Agricultural Institute (Menangle, New South Wales, Australia) from semi-purified liver homogenates after amplification in domestic rabbits, as described previously [1, 32]. These virus stocks were titrated *in vivo* to determine the 50% rabbit infectious dose (RID_50_). Freeze-dried virus stocks were reconstituted in sterile water and stored at −80°C between experiments.

### Inoculation of hepatobiliary organoid monolayer cultures

One-day-old confluent monolayer cultures grown in 96-well plates or 8-well chamber slides were inoculated in triplicate with either RHDV1, RHDVa-K5, or RHDV2 diluted in culture medium without antibiotic/antimycotic in a total volume of 100 μl or 200 μl, respectively. We used an infectious dose of 100 RID_50_ per well for 96-well plates or 200 RID_50_ per chamber for chamber slides. This equates to a multiplicity of infection (MOI) of approximately 0.001 (based on 7 × 10^4^ cells per well for 96-well plates and 1.4 × 10^5^ cells per chamber for chamber slides), and to a log_10_ capsid gene copy number (per 100 μl) of 8.8 (RHDV1), 6.7 (RHDVa-K5) and 5.3 (RHDV2), based on RT-qPCR quantification of each inoculum preparation (as described below). Inoculum that had been heat-inactivated at 63°C for 15 minutes was used as a control. Virions were adsorbed for 1 hour at 37°C in 5% CO_2_. For each inoculation experiment, a baseline plate, referred to as 1-hour post inoculation (hpi), was harvested after this adsorption step. For other time points, the inoculum was removed and replaced with fresh culture medium (100 μl for 96-well plates or 200 μl for chamber slides) and plates were harvested at specific timepoints, as detailed for each experiment. For experiments where immunofluorescence staining was conducted, chamber slides were infected in parallel and fixed in 4% paraformaldehyde at the required timepoints after washing in PBS.

### RNA extraction and RT-qPCR

Monolayer cultures were harvested from 96-well plates by first collecting the culture supernatant and then harvesting cells with a cell scraper and adding these to the supernatant. Total RNA was isolated (from the entire contents of the well, i.e., cells and supernatant) using either the NucleoSpin RNA II Mini kit (Macherey-Nagel) or the RNeasy Mini kit (Qiagen), as per manufacturers’ instructions. For gene expression analyses, organoids were harvested using TrypLE Express Enzyme (Thermo Fisher Scientific) and pelleted at 200 *g* for 5 minutes at 4°C. Total RNA was extracted from cell pellets using the NucleoSpin RNA II Mini kit (Macherey-Nagel), as per manufacturer’s instructions. Total RNA was isolated from liver tissue collected from a healthy, adult New Zealand White laboratory rabbit (CWLA-AEC 2016-22). For quantification of inocula, RNA was extracted from 50 μl of virus inoculum suspension using the PureLink Viral RNA/DNA Mini kit (Themo Fisher Scientific), as per manufacturer’s directions.

For gene expression analyses, RT-qPCR was performed using the SensiFAST SYBR No-ROX kit (Bioline) according to the manufacturer’s instructions on a CFX96 Touch real-time PCR instrument (Bio-Rad). Primers for gene expression analyses were designed to be intron spanning and between 17–21 bases in length using NCBI Primer-BLAST (Supplementary Table 3). The following cycling conditions were used: reverse transcription at 45°C for 10 minutes, followed by initial denaturation at 95°C for 5 minutes, then 40 cycles of 95°C for 10 seconds, 63°C for 40 seconds and 78°C for 10 seconds, with a melt curve analysis from 95°C to 65°C in 0.5°C increments for 5 seconds. ‘No template’ and ‘no reverse transcriptase’ controls were performed with each run. Transcription of the gene of interest was normalised to expression levels of a housekeeping gene (GAPDH) (Supplementary Table 3). Fold change was calculated using the 2^-ΔΔCt^ method.

For infection experiments, viral RNA was quantified using a previously described universal RT-qPCR assay targeting a conserved region of lagovirus VP60 and utilising a standard curve for absolute quantification [33]. Results are reported in units of log_10_ capsid gene copies per well. For analysis purposes, samples where no viral RNA was detected were set to 1 capsid gene copy per well. A sample was considered positive where template was detected in both technical replicates.

### Immunofluorescence staining

For immunofluorescence staining, monolayer cultures grown in chamber slides were washed twice with PBS and fixed in 4% paraformaldehyde in PBS for 15 minutes at room temperature, then washed again twice with PBS. Citrate buffer (10 mM) was added and the slides were incubated for 20 minutes at 95°C to expose antigenic sites. Following antigen retrieval, the slides were cooled and washed with PBS for 5 minutes. The slides were then permeabilised with 0.25% Triton X-100 in PBS for 10 minutes at room temperature, washed with PBS for 5 minutes and blocked with 5% bovine serum albumin (BSA) in PBS with 0.25% Tween 20 for 20 minutes. The following antibodies were used: the lagovirus-specific anti-VP60 (capsid) mouse monoclonal antibody 13C10 (25 μg/ml; Monoclonal Antibody Facility of the Institute for Medical Research, Perth, Australia) [34], the lagovirus-specific anti-polymerase mouse monoclonal antibody 8G9 (Walter and Eliza Hall Institute of Medical Research Antibody Facility, Melbourne, Australia), a double-stranded RNA antibody (dsRNA; 10 μg/ml; Scicons) and the cytokeratin-19 antibody (KRT-19; Abcam; for further details, see Supplementary Table 2). All primary antibodies were diluted in PBS containing 0.25% Tween 20 and 1% BSA, and slides were placed into a humidified chamber and incubated overnight at 4°C. Slides were then washed two times with PBS for 5 minutes before the appropriate secondary antibodies, diluted in PBS with 1% BSA, were added. The slides were incubated at room temperature for another 40 minutes, washed two times with PBS, counter-stained with 4’,6-diamidino-2-phenylindole (DAPI) for 5 minutes at room temperature and mounted with Fluoroshield (Sigma-Aldrich).

### Statistical analyses

Statistical differences in gene expression were assessed using one-way analysis of variance (ANOVA) followed by the Tukey’s honestly significant difference test (Tukey’s HSD), as calculated using the BioRad CFX Maestro software (BioRad). Statistical significance was assessed at *p* = 0.05.

## Results

### Characterisation of rabbit hepatobiliary organoids and organoid-derived monolayer cultures

To assess whether rabbit hepatobiliary organoids could support lagovirus replication, we first generated 3D cultures from rabbit liver biopsies. Hepatobiliary organoids were expanded for up to 15 passages in a human liver organoid culture medium reported previously [28], with minor modifications (Figure 2A). These organoids were able to be transformed into 2D monolayer cultures and could be cryopreserved and revived without any morphological changes.

**Figure 2.**
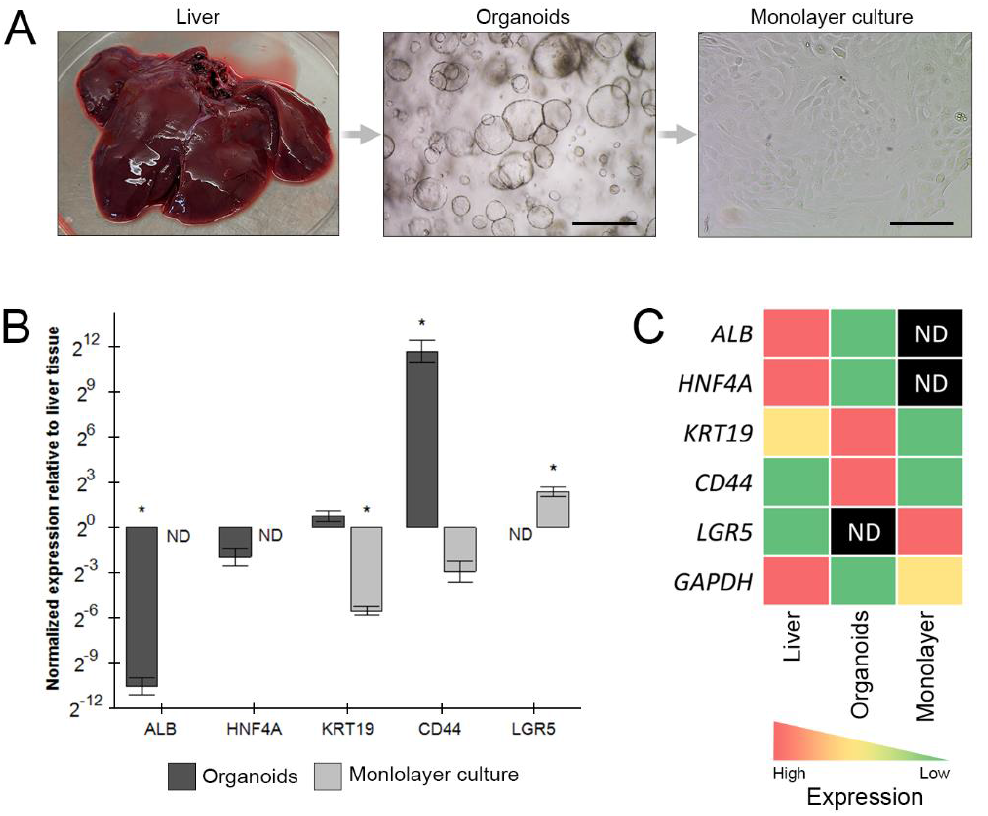
Characterisation of rabbit hepatobiliary organoids and organoid-derived monolayer cultures. (A) Hepatic progenitor cells were isolated from rabbit liver tissue (left) and propagated in culture medium to generate 3D organoids (middle). Organoids could be dissociated into 2D monolayer cultures (right). Scale bars = 500 μm. (B) Expression of hepatocyte-associated (*albumin* (*ALB*), *hepatocyte nuclear factor-4 alpha* (*HNF4A*)) and cholangiocyte-associated (*cytokeratin-19* (*KRT19*)) genes in 3D rabbit hepatobiliary organoids and 2D organoid-derived monolayer cultures, relative to rabbit liver tissue. Also shown are the epithelial cell marker *cluster of differentiation marker 44* (*CD44*) and the stem cell marker *leucine-rich repeat-containing G-protein coupled receptor 5* (*LGR5*). Data are presented as the fold change (2^-ΔΔCt^) in gene expression using *GAPDH* as the housekeeping gene. Means were calculated from three independent biological replicates with two technical replicates each. Error bars represent the standard error of the mean. Asterisks indicate statistical significance (*p* <0.05), as determined by one-way analysis of variance (ANOVA) followed by the Tukey’s honest significant difference test (Tukey’s HSD). (C) Heatmap of expression levels of *ALB, HNF4A, KRT19, CD44* and *LGR5* in rabbit liver tissue, hepatobiliary organoids and organoid-derived monolayer cultures. ND, not detected.

We then characterised the cell populations of these organoids and monolayer cultures in comparison to native liver tissue using gene expression analyses. We looked at expression levels for albumin (*ALB;* a mature hepatocyte marker), hepatocyte nuclear factor-4 alpha (*HNF4A;* expressed in a wide range of cells, including hepatoblasts and hepatocytes, but downregulated in cholangiocytes [35]), leucine-rich repeat-containing G-protein coupled receptor 5 (*LGR5;* a stem cell marker [30]), KRT-19 (a marker of cholangiocytes [36]) and cluster of differentiation marker 44 (*CD44;* an epithelial cell marker), and used the housekeeping gene glyceraldehyde-3-phosphate dehydrogenase (*GAPDH*) for normalisation. As expected, liver tissue expressed high levels of *ALB* and *HNF4A*, moderate levels of *KRT19*, and lower levels of *LGR5* and *CD44* (Figure 2B, 2C). In contrast, hepatobiliary organoids showed high expression of *KRT19* and *CD44* and low expression of the hepatocyte markers *ALB* and *HNF4A*, suggestive of a cholangiocyte phenotype, and no *LGR5* expression was detected (Figure 2B, 2C). The pattern of gene expression in organoid-derived monolayer cultures was similar to that observed in organoids, with the exception of *LGR5*, which was expressed at high levels (Figure 2B, 2C). While *ALB* and *HNF4A* were not detected in the monolayer cultures, the quantity of RNA obtained was very low, as shown by the low expression levels of *GAPDH* relative to organoids and liver tissue (Figure 2C). Taken together, these results demonstrate that our rabbit hepatobiliary organoids and organoid-derived monolayer cultures contain a predominantly biliary cell population but may lack mature hepatocytes.

### Only hepatobiliary organoids derived from leporids support lagovirus replication

To explore the biological relevance of our rabbit organoids, we additionally generated hepatobiliary organoids from a European brown hare (*Lepus europaeus*), a wild mouse (*Mus musculus*) and a feral cat (*Felis domesticus*) and infected organoid-derived monolayer cultures with three genotypes of lagovirus: GI.1cP-GI.1c (RHDV1), GI.1aP-GI.1a (RHDVa-K5) and GI.1bP-GI.2 (RHDV2). While RHDV2 can infect rabbits (*Orycotolagus*), hares (*Lepus),=* and cottontails (*Sylvilagus*) [5–12], RHDV1 and RHDVa viruses are generally considered to infect only rabbits [2, 37]. We quantified the change in viral capsid gene copies between 1 hpi, i.e., after the 1-hour adsorption period, and 24 hpi as a measure of viral replication. We observed a consistent increase of greater than one log_10_ capsid gene copies between 1 and 24 hpi across all three biological replicates of rabbit monolayer cultures infected with RHDV1, RHDVa-K5 and RHDV2, and for hare monolayer cultures infected with RHDV2 only (Figure 3A, Table 1). While RT-qPCR is a highly sensitive technique, it is imperative to differentiate between limit of detection (LoD) and the limit of quantitation (LoQ) [38]. The LoD is the analytical sensitivity or the lowest amount of target that can be detected; however, particularly at low target concentrations, there can be considerable imprecision or “noise” that is further compounded by the risk of aerosol contamination from positive samples and controls. In contrast, the LoQ is the lowest amount of target that can be quantitatively determined with acceptable precision and accuracy. The LoQ for the assay used in this study was previously determined to be is 2 log_10_ capsid copies per μl RNA [33]. While we also observed a greater than one log_10_ increase in cat hepatobiliary monolayer cultures infected with RHDV2 and uninfected cat and rabbit monolayer cultures, this increase was not consistent across replicates and the mean increase in viral RNA levels was near or below the LoQ of the assay (Figure 3A, Table 1). When virus inocula were heat-inactivated, a maximum increase of 0.3 log_10_ capsid gene copies per well between 1 and 24 hpi was observed (Table 1). Interestingly, while all monolayer cultures were infected with 100 RID_50_ per well and a single mastermix of inoculum was prepared for each virus, the virus RNA levels at 1 hpi was at least one log_10_ capsid gene copies higher in rabbit cultures for all three viruses and in hare cultures for RHDVa-K5 and RHDV2 (Figure 3A), perhaps suggesting early replication that preceded the baseline 1 hpi timepoint or enhanced adsorption of the respective viruses to leporid cells. The RNA levels for heat-inactivated virus at 1 hpi was also lower than for the corresponding non-heat-treated inocula, indicating that heat-treatment likely destroyed some viral RNA despite being it protected within the capsid.

**Figure 3.**
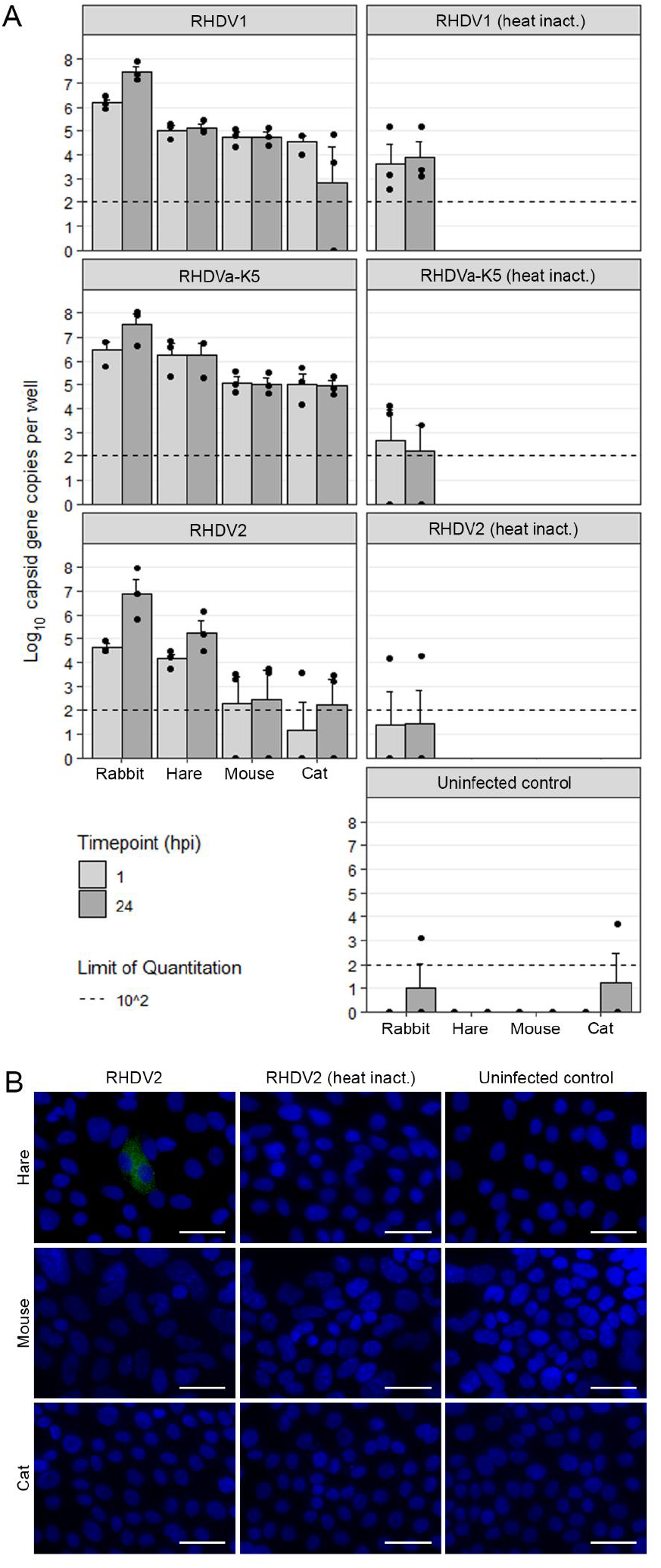
Inoculation of hepatobiliary organoid-derived monolayer cultures derived from rabbit, hare, mouse and cat with various lagoviruses. Monolayer cultures derived from rabbit, hare, mouse and cat hepatobiliary organoids were inoculated with 100 ‘50% rabbit infectious doses’ (RID_50_) of either RHDV1, RHDVa-K5, or RHDV2 lagoviruses. Heat-inactivated inocula and uninfected monolayer cultures were used as controls. Monolayer cultures were analysed at 1 and 24 hours post inoculation (hpi). (A) Viral RNA per well was quantified at 1 and 24 hpi using RT-qPCR. The mean of three biological replicates is shown as a bar and the individual data points are shown at each time point (filled circles). Error bars represent the standard error of the mean. The limit of quantitation of the assay is shown by a horizontal line. (B) Monolayer cultures derived from hare, mouse and cat hepatobiliary organoids were grown in chamber slides and inoculated with 200 RID_50_ of RHDV2 (to maintain the same multiplicity of infection), as described above. Cultures were immunostained at 24 hpi with a mouse monoclonal antibody directed against the RHDV2 RNA-dependent RNA polymerase (RdRp; green); DNA was stained with DAPI (blue). Scale bars = 50 μm.

**Table 1.** Mean log_10_ change in capsid gene copies per well between 1 and 24 hours post inoculation. Rabbit, hare, mouse and cat hepatobiliary organoid-derived monolayer cultures were inoculated with various lagoviruses. Means were calculated based on three biological replicates with two technical replicates each. ND, not done.

| Inoculum virus | Rabbit | Hare | Mouse | Cat |
| --- | --- | --- | --- | --- |
| RHDV1 | 1.31 | 0.11 | -0.00 | -1.68 |
| RHDVa-K5 | 1.09 | 0.00 | -0.06 | -0.06 |
| RHDV2 | 2.24 | 1.12 | 0.16 | 1.04 |
| Heat-inactivated RHDV1 | 0.26 | ND | ND | ND |
| Heat-inactivated RHDVa-K5 | -0.43 | ND | ND | ND |
| Heat-inactivated RHDV2 | 0.04 | ND | ND | ND |
| Uninfected | 1.03 | 0 | 0 | 1.23 |

In addition to the RT-qPCR analysis, we further assess the species tropism of RHDV2, we inoculated hare, mouse and cat hepatobiliary organoid-derived monolayer cultures with either RHDV2 or heat-inactivated RHDV2 and looked for expression of the viral RNA-dependent RNA polymerase (RdRp) by immunofluorescence. The viral RdRp is a non-structural protein and as such, would not be present in virus particles but must be expressed after infection. We observed immunofluorescence staining only in hare monolayer cultures infected with RHDV2, further suggesting that hare, but not mouse or cat, monolayer cultures are permissive for RHDV2 infection (Figure 3B). Taken together, our results demonstrate that hepatobiliary organoid-derived monolayer cultures (*i*) support the replication of hepatotropic lagoviruses, and (*ii*) simulate the *in vivo* species-specificity of these viruses.

### Time course of RHDV2 replication in rabbit hepatobiliary organoids

To investigate the kinetics of viral replication, we infected rabbit hepatobiliary-derived monolayer cultures with 100 RID_50_ of RHDV2 and looked for viral RNA by RT-qPCR and for expression of the lagovirus VP60 (capsid) protein and double-stranded RNA (dsRNA; a canonical viral replication intermediate) by immunofluorescence staining at 1, 24, 48, 72 and 96 hpi. We observed an increase in viral RNA until 48 hpi, followed by a gradual decline to 96 hpi (Figure 4A), while the detection of lagovirus VP60 and dsRNA peaked at 24 hpi (Figure 4B). From 48 hpi, cells and cellular debris stained positively for either VP60 or dsRNA only sporadically. Interestingly, when infected monolayer cultures were co-stained for VP60 and KRT19, most infected cells had a KRT19^-^ phenotype (Figure 4B), suggesting that cholangiocytes are not a primary virus target in hepatobiliary organoid cultures. Infected monolayer cultures appeared similar to uninfected controls by brightfield microscopy, and no cytopathic effect was observed. The consistent detection of viral RNA, viral capsid protein and dsRNA in rabbit hepatobiliary-derived monolayer cultures conclusively demonstrates that this is a robust and reliable *ex vivo* model for lagovirus infection.

**Figure 4.**
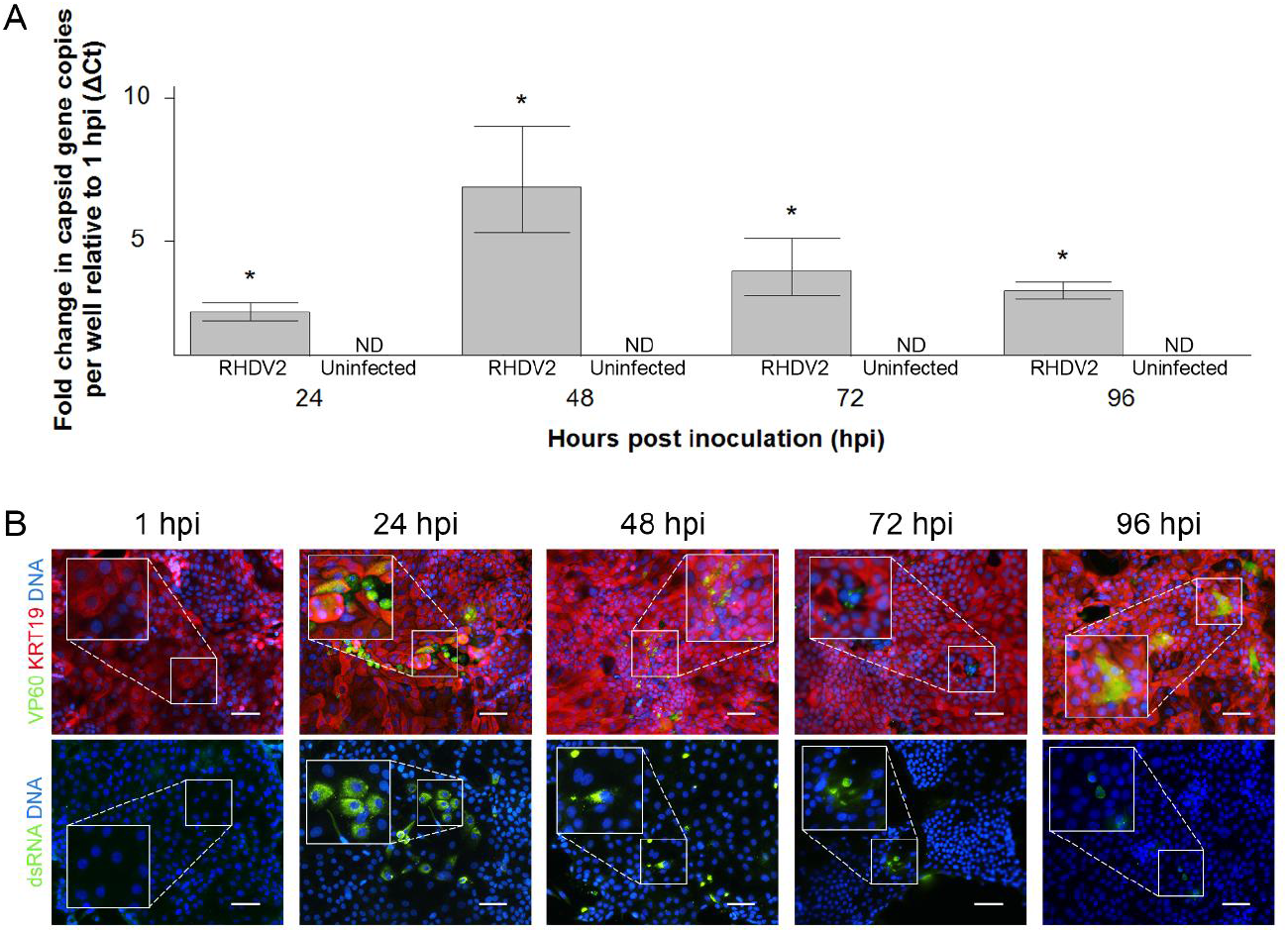
Kinetics of RHDV2 replication in rabbit hepatobiliary organoid-derived monolayer cultures. Monolayer cultures derived from rabbit hepatobiliary organoids were inoculated with 100 ‘50% rabbit infectious doses’ (RID_50_) of RHDV2 and were harvested at 1, 24, 48, 72 and 96 hours post inoculation (hpi). Uninfected monolayer cultures were run as controls. (A) Viral RNA per well was quantified at each time-point using RT-qPCR. The mean fold change compared to 1 hpi at each time-point is plotted, based on three biological replicates. Error bars represent the standard error of the mean. Significance (relative to 1 hpi), denoted by an asterisk, was assessed using one-way analysis of variance (ANOVA) followed by Tukey’s honest significance test, with a *p*-value cut-off of 0.05. (B) Monolayer cultures grown in chamber slides were inoculated as described above and immunostained at each timepoint with monoclonal antibodies directed against the lagovirus capsid protein (VP60; green), cytokeratin-19 (KRT19; red), or double-stranded RNA (dsRNA; green). DNA was stained with DAPI (blue). Scale bars = 100 μm. ND, not detected.

## Discussion

Despite extensive efforts spanning several decades, lagoviruses have long resisted *ex vivo* cultivation in all systems apart from primary rabbit hepatocytes [2]. Here we report the development of a robust *ex vivo* model for lagoviruses using hepatobiliary organoid-derived monolayer cultures generated from rabbit and hare liver samples. Unlike primary hepatocytes, hepatobiliary organoids are self-renewing, which enables indefinite propagation and expansion of these cells from cryopreserved stocks [23]. Indeed, we were able to maintain our rabbit hepatobiliary organoids for at least 15 passages.

While there is clearly more work needed to further characterise and optimise this system, we conclusively demonstrated that our leporid hepatobiliary organoid-derived monolayer cultures support the replication of RHDV2. We consistently observed a greater than one log_10_ increase in viral RNA copies per infected well over a 23-hour period; we immunostained double-stranded RNA (a canonical viral replication intermediate), suggesting active viral replication; and we detected expression of the viral capsid protein and the non-structural RdRp in infected leporid monolayer cultures.

Contrastingly, we did not observe antigen expression after inoculation with heat-inactivated virus or in cells from non-permissive animals. We also used Northern blot assays to detect negative-sense viral RNA in infected cultures, but the relatively low sensitivity of this assay did not allow us to detect the moderate viral RNA loads obtained in our 96-well plate monolayer culture system (Supplementary Figure 1) [39]. We found considerable variability in viral RNA levels both between replicates within an experiment and between experiments; however, this inherent variability is a widely recognised characteristic of organoid culture systems [40, 41]. For example, replication of a GII.4 human norovirus in intestinal organoid-derived monolayer cultures over a similar 23-hour period was shown to vary from just over one log_10_ to 3.38 log_10_ genome equivalents per well, based on 80 independent experiments over a 3-year period with considerable variability between replicates [40]. The large variance and small sample sizes (n = 3 replicates) in our study limited the statistical analysis. Without confirmatory immunostaining, we recognise that the evidence for replication of RHDV1 and RHDVa-K5 in rabbit monolayer cultures may be considered circumstantial.

Our rabbit organoid and monolayer cultures were heavily biased towards the cholangiocyte phenotype; negligible to very low expression levels of *ALB* and *HNF4A* were detected. This phenotype appears to be a common (and a known issue) for hepatobiliary organoids [30, 42]. We did attempt to differentiate our monolayer cultures to drive differentiation towards a mature hepatocyte fate; however, we used a 15-day differentiation protocol [28], over which time many of the cells died. Based on co-immunostaining for the RHDV2 capsid protein and the cholangiocyte marker KRT-19, most of the viral antigens were detected in cells with a KRT-19-negative phenotype. Importantly, the liver consists largely of hepatocytes [19, 20] and hepatocytes are known to be the primary target cells of lagovirus infection *in vivo* [3]. Therefore, future work to manipulate the differentiation pathways of our rabbit hepatobiliary organoids to better represent the native rabbit liver tissue will be required to optimise this *ex vivo* model.

To standardise infection experiments with different viruses we consistently used 100 RID_50_, since infectious dose has been shown to affect disease progression *in vivo* [1]. Titrations of viral stocks were done in rabbits [1, 32] but it is unknown how well rabbit infectious doses correlate with cell culture infectious doses. Nevertheless, we considered RID_50_ to be a better measure of viral infectivity than capsid gene copy numbers. However, it is worth noting that capsid gene copy numbers per 100 RID_50_ varied widely between the different lagovirus variants tested. We assumed that measuring the change in viral RNA levels between 1 and 24 hpi would counteract these differences between variants, however, for future studies it will be important to establish a reliable focus-forming assay (since cytopathic effects were not observed) to better standardise between variants and to correlate *in vivo* with *ex vivo* infectivity and capsid gene copies. Conspicuously, RHDV1 and RHDVa-K5 had much higher capsid gene copies per 100 RID_50_ than did RHDV2, potentially limiting the observable increase in viral RNA for these variants. Notably, long-term passaging of human norovirus and the generation of high-titre viral stocks using organoids has previously proven challenging [41]. Whether these limitations also exist for lagoviruses is unclear, as we did not attempt to passage viruses in this study.

Several studies have previously claimed that lagoviruses can infect non-leporid species, including wood mice (*Apodemus sylvaticus*), voles (*Microtus duodecomcostatus*), shrews (*Crocidura russula*) and badgers (*Meles meles*) [43–46]. It is well-established that antibodies against lagoviruses and/or trace amounts of viral genomic RNA can be detected in non-leporid species, especially predators and scavengers like foxes, dogs and cats [47–50]. However, definitive evidence of active (non-abortive) viral replication, such as an increase in viral RNA levels over time, the presence of viral non-structural antigens in target tissues in association with histopathological changes, and/or the presence of replicative intermediates, in non-leporids has not been demonstrated. Furthermore, experimental studies where cats were fed RHDV1-infected livers failed to demonstrate active replication [51]. Even in immunocompromised mice that lack a functional interferon type I receptor, challenge studies did not reveal evidence of virus amplification [34]. To investigate lagovirus species specificity using our organoid model, we grew hepatobiliary organoids from several non-rabbit species (hare, mouse and cat) and infected them with several lagovirus variants. We showed that the hepatobiliary organoids recapitulate the species tropism observed for the different variants *in vivo*, namely that replication of RHDV1 and RHDVa-K5 is restricted to rabbits, while RHDV2 is capable of infecting both rabbit and hare cells. The organoid cultures from non-leporid species (cat and mouse) did not support lagovirus replication. Our results support the notion that previous claims of lagovirus infection in non-leporid species are likely due to detection of trace amounts of viral RNA after ingestion of large amounts of virus (in the absence of productive infection) or may be PCR artefacts.

To conclude, we have established hepatobiliary organoids and organoid-derived monolayers cultures from rabbit, hare, cat and mouse liver tissue. Monolayer cultures from rabbit and hare, but not cat or mouse, supported replication of RHDV2, as evidenced through an increase in viral RNA levels over time, expression of viral structural and non-structural proteins detected by immunostaining, and the presence of double-stranded RNA in infected leporid-derived cultures. In addition to greatly facilitating studies into the fundamental biology of these hypervirulent and important viruses (in their context as biocontrol agents to manage wild rabbit populations in Australia), this *ex vivo* model will help to reduce the reliance on animals for many aspects of lagovirus research [23]. The addition of the rabbit organoid model to the suite of existing lagovirus research models, namely *in vitro* assays, *in vivo* rabbit infection models, and extensive field data accumulated over many years and across continents, presents a unique opportunity to disentangle the mechanisms of virulence and tropism of caliciviruses more broadly.

## Author Statements

### Authors and contributors

Conceptualisation – RNH, MF, TS, ME; Methodology – EK, RNH, MF, TS, ME; Investigation – EK, OF, MP, HM, NH, ES; Formal analysis – OF, HM, RNH; Data curation – EK, RNH, OF. MP; Writing – Original draft preparation – EK, RNH; Writing – Review and editing – EK, OF, MP, HM, NH, ES, ME, TS, MF, RNH; Visualisation – OF, EK, RNH; Supervision – RNH, MF, TS; Project administration – RNH; Funding – TS, RNH.

### Ethics approval

All animal use in this study was approved by the CSIRO Wildlife and Large Animal - Animal Ethics Committee (CWLA-AEC permit numbers 2018-06, 2019-28, 2020-24 and 2020-01). All animal procedures were carried out at the CSIRO Black Mountain Laboratories according to the Australian Code for the Care and Use of Animals for Scientific Purposes and in compliance with the ARRIVE guidelines.

## Acknowledgements

We wish to acknowledge the following colleagues for assistance with sample collection: Melissa Piper (CSIRO Land & Water) for domestic rabbit samples; Georgeanna Story (Upper Murrumbidgee Landcare Committee) and Alison and Richard Swain (Alpine River Adventures) for the feral cat liver sample; Kevin Oh (Macquarie University Applied BioSciences and CSIRO Health & Biosecurity) for the wild mouse liver sample; and Amelia Keyworth (ACT Parks and Conservation Service) for the wild hare liver sample. We thank Stewart Nuttall and Anna Raicevic (CSIRO Manufacturing) for assistance with expression of the RHDV2 polymerase for monoclonal antibody production. We thank Sarron Randall-Demllo, Theo Almond and Kim Fung (CSIRO Health & Biosecurity) for thoughtful discussions. We also thank Kim Fung and Helen Palethorpe for critical revision of the manuscript.

## Supplementary materials

**Supplementary Table 1.** Media formulations and reagents used in this study, adapted from those in Broutier et al [28].

| Number | Component | Supplier | Catalogue number | Working concentration |
| --- | --- | --- | --- | --- |
| 1 | Basal medium – 1-month storage life at 4°C |  |  |  |
|  | Advanced Dulbecco's modified Eagle medium (DMEM)/F-12 | Thermo Fisher Scientific | 12634-010 |  |
|  | Antibiotic/antimycotic | Thermo Fisher Scientific | 15240096 | 1 x |
|  | GlutaMAX | Thermo Fisher Scientific | 35050-061 | 2 mM |
|  | N-2-hydroxyethylpiperazine-N'-2-ethanesulfonic acid (HEPES) | Thermo Fisher Scientific | 15630-056 | 10 mM |
| 2 | Wash medium |  |  |  |
|  | 1x phosphate-buffered saline (PBS) |  |  |  |
| 3 | Digestion medium – prepare fresh |  |  |  |
|  | Earle's balanced salt solution (EBSS) | Thermo Fisher Scientific | 24010043 |  |
|  | Collagenase D | Roche | 11088858001 | 2.5 mg/ml |
|  | DNaseI | Sigma-Aldrich | DN25 | 0.1 mg/ml |
| 4 | Culture medium – 2-week storage life at 4°C |  |  |  |
|  | Basal medium (#1) |  |  |  |
|  | B27 supplement | Thermo Fisher Scientific | 17504-044 | 1:50 |
|  | N2 supplement | Thermo Fisher Scientific | 17502-048 | 1:100 |
|  | N-acetylcysteine | Sigma-Aldrich | A9165 | 1mM |
|  | Nicotinamide | Sigma-Aldrich | N0636 | 10 mM |
|  | Human [Leu15]-gastrin I | Sigma-Aldrich | G9145 | 10 nM |
|  | Human epidermal growth factor (EGF) | Invitrogen | PHG0311 | 50 ng/ml |
|  | Human fibroblast growth factor (FGF) 10 | Stemcell Technologies | 78037 | 60 ng/ml |
|  | Human hepatocyte growth factor (HGF) | Stemcell Technologies | 78019.1 | 25 ng/ml |
| | Forskolin | Cayman Chemicals | 11018 | 10 μM |
| | A83-01 (TGF-β inhibitor) | Sigma-Aldrich | SML0788 | 5 μM |
|  | L-WRN conditioned media | Prepared as described<br>previously [31] |  | 55% |
| | Y-27632 (rho kinase [ROCK] inhibitor) | Cayman Chemicals | 129830-38-2 | 10 μM |
| 5 | Additional reagents |  |  |  |
|  | Freezing solution | Thermo Fisher Scientific | 12648-010 |  |
|  | TrypLE Express Enzyme | Thermo Fisher Scientific | 12604021 |  |
|  | 0.25% Trypsin - ethylenediaminetetraacetic acid (EDTA) | Thermo Fisher Scientific | 25200072 |  |
|  | Geltrex lactose dehydrogenase elevating virus free (LDEV-free)<br>reduced growth factor basement membrane matrix | Thermo Fisher Scientific | A1413202 | 1:50 |

**Supplementary Table 2.** Antibodies used in this study.

| **Antibody name** | **Specificity** | **Product number** | **Supplier** | **Conjugation** |
| --- | --- | --- | --- | --- |
| Mouse monoclonal lagovirus-specific anti-VP60 (capsid) | Lagovirus VP60 (capsid) protein [34] | Mab-13C10 | Monoclonal Antibody Facility of the Institute for Medical Research, Perth, Australia | Unconjugated |
| Mouse monoclonal lagovirus-specific anti-polymerase | Lagovirus RNA-dependent RNA polymerase | Mab-8G9 | Walter and Eliza Hall Institute of Medical Research Antibody Facility, Melbourne, Australia | Unconjugated |
| Mouse monoclonal anti-dsRNA | Double-stranded RNA | J2-1402 | Scicons | Unconjugated |
| Rabbit monoclonal anti-Cytokeratin 19 (KRT-19) | Keratins of epithelial cells; biliary/progenitor cell marker in the liver | ab76539 | Abcam | Unconjugated |
| Secondary goat anti-mouse | Goat polyclonal secondary antibody to mouse IgG | ab150113 | Abcam | Alexa Fluor 488 |
| Secondary goat anti-mouse | Goat polyclonal secondary antibody to mouse IgG | ab150114 | Abcam | Alexa Fluor 555 |
| Secondary goat anti-rabbit | Goat polyclonal secondary antibody to rabbit IgG | ab150078 | Abcam | Alexa Fluor 555 |

**Supplementary Table 3.**
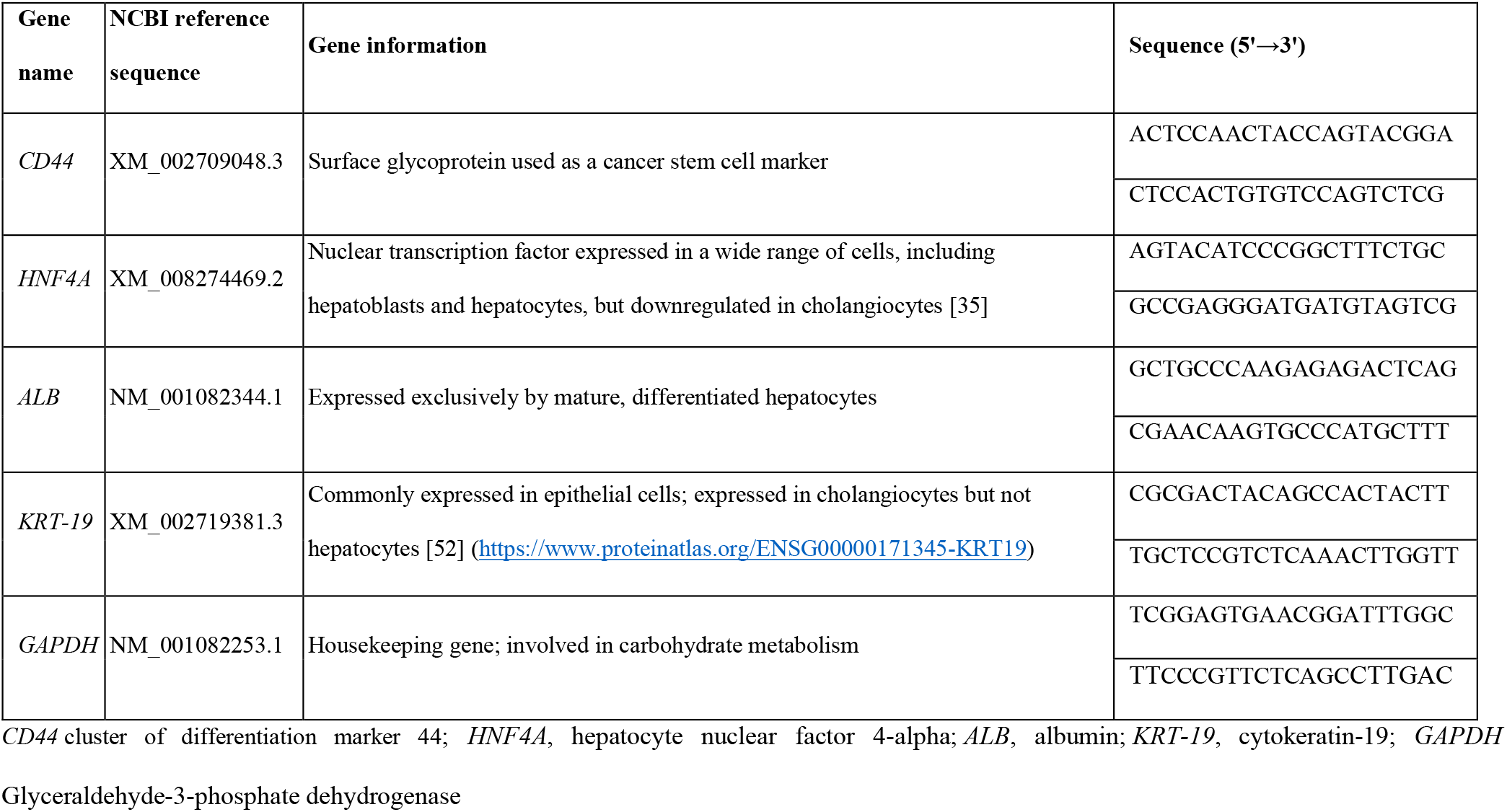
Primers used in this study.

**Supplementary Figure 1.**
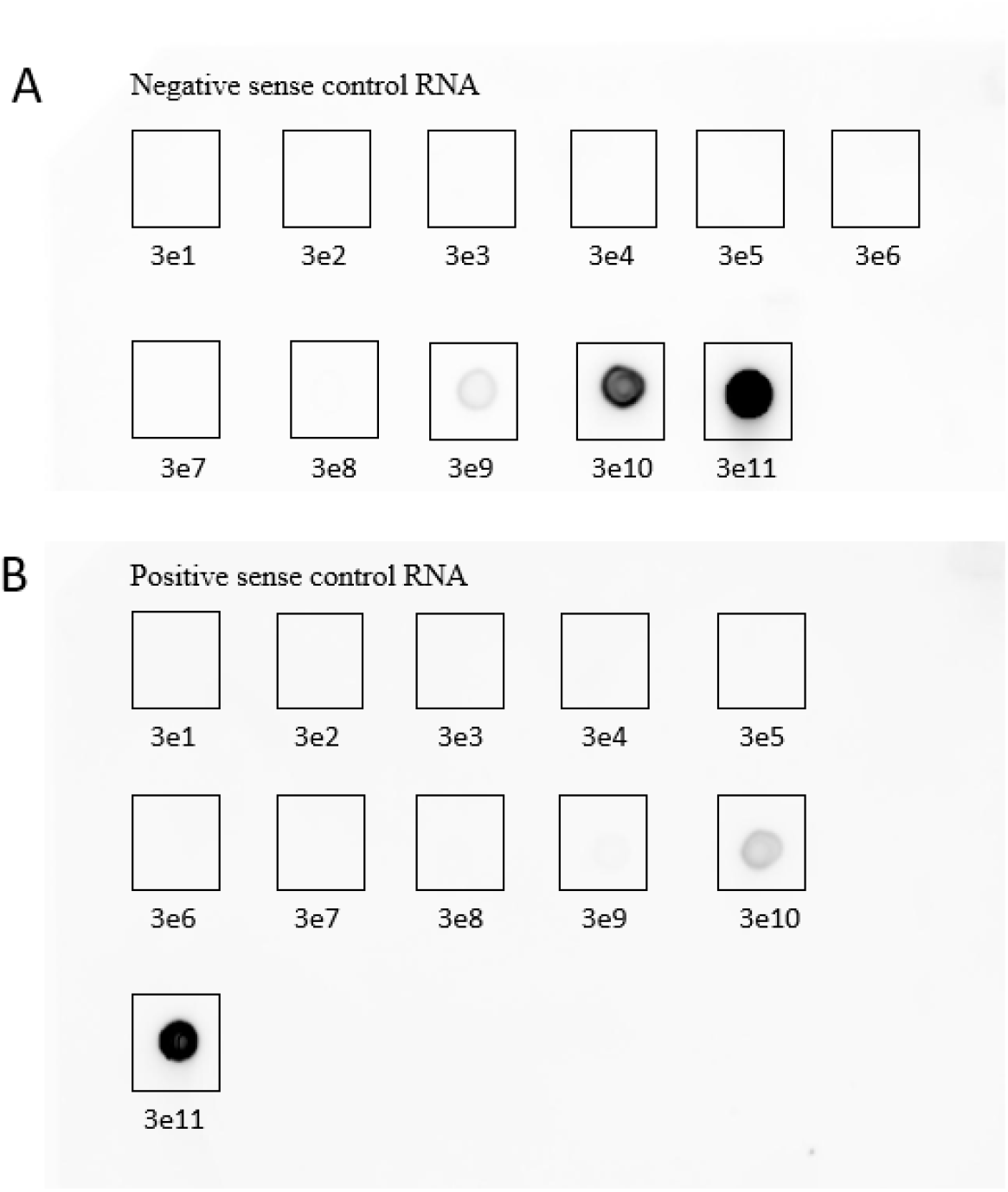
Sensitivity testing of a lagovirus-specific Northern blot assay. To generate positive control RNA transcripts to assess the sensitivity of our Northern blot assay, primers were designed that amplified a 561-bp region within the RHDV2 VP60 (capsid) gene (GenBank accession MW467791). This region was selected to maximise sensitivity because the target is also present within the colinear viral subgenomic RNA. Two sets of primers were designed, to generate a positive sense control template and probe and a negative sense control template and probe, respectively. For each set the forward primer was tailed at the 5’ end with a T7 promoter sequence (denoted in bold) to enable *in vitro* transcription (positive sense forward: 5’-**TAATACGACTCACTATAGGA**GTGGCATGCAATTCCGGTTUAU-3’; positive sense reverse: 5-CGUUACCUUUGUCGUGGUCAAU-3’; negative sense forward: 5’-**TAATACGACTCACTATAGGA**CGUUACCUUUGUCGUGGUCAAU-3’; negative sense reverse: 5’-GUGGCAUGCAAUUCCGGUUUAU-3’). Viral RNA was extracted from the RHDV2 virus inoculum as described in the Methods and template cDNA was generated using SuperScript IV reverse transcriptase and 500 ng of oligodT primer (Thermo Fisher Scientific), according to the manufacturer’s directions. The cDNA was amplified with the primers described above (at a final concentration of 300 nM) using Platinum Taq DNA polymerase (Thermo Fisher Scientific) with the following cycling conditions: 94°C for 2 minutes, followed by 30 cycles of 94°C for 45 seconds, 60°C for 45 seconds and 72°C for 90 seconds, with a final extension at 72°C for 10 minutes. The amplicons were used as template to generate *in vitro* transcript controls with the SP6/T7 Riboprobe Combination System (Promega), as per manufacturer’s instructions. Transcripts were quantified with the Qubit RNA HS assay (Thermo Fisher Scientific) from which we calculated the copy number per μl of RNA. A ten-fold dilution series from 10^11^ to 10^1^ copies per μl was prepared for each transcript. Digoxigenin (DIG)-labelled RNA hybridisation probes were generated using the DIG Northern Starter Kit (Roche); probes were quantified Qubit RNA HS assay (Thermo Fisher Scientific). Control transcripts (3 μl per dilution; negative sense (A) and positive sense (B)) were spotted onto a positively-charged nylon membrane (Sigma-Aldrich) and cross-linking was performed by incubating at 80°C for 20 minutes. Hybridisation was performed with the respective complementary DIG-labelled probe using the DIG Northern Starter Kit (Roche) as per manufacturer’s directions. Membranes were imaged using an Amersham Imager 600 (GE Healthcare).

## References

1. Hall RN, King T, O’Connor T, Read AJ, Arrow J et al. Age and infectious dose significantly affect disease progression after RHDV2 infection in naive domestic rabbits. Viruses 2021;13(6) 10.3390/v13061184.

2. Abrantes J, van der Loo W, Le Pendu J, Esteves PJ. Rabbit haemorrhagic disease (RHD) and rabbit haemorrhagic disease virus (RHDV): a review. Veterinary Research 2012;43:12 10.1186/1297-9716-43-12.

3. Neimanis A, Larsson Pettersson U, Huang N, Gavier-Widen D, Strive T. Elucidation of the pathology and tissue distribution of *Lagovirus europaeus* GI.2/RHDV2 (rabbit haemorrhagic disease virus 2) in young and adult rabbits (*Oryctolagus cuniculus*). Veterinary Research 2018;49(1):46 10.1186/s13567-018-0540-z.

4. Le Pendu J, Abrantes J, Bertagnoli S, Guitton JS, Le Gall-Recule G et al. Proposal for a unified classification system and nomenclature of lagoviruses. Journal of General Virology 2017;98(7):1658–1666 10.1099/jgv.0.000840.

5. Asin J, Nyaoke AC, Moore JD, Gonzalez-Astudillo V, Clifford DL et al. Outbreak of rabbit hemorrhagic disease virus 2 in the southwestern United States: first detections in southern California. Journal of Veterinary Diagnostic Investigation 2021;33(4):728–731 10.1177/10406387211006353.

6. Camarda A, Pugliese N, Cavadini P, Circella E, Capucci L et al. Detection of the new emerging rabbit haemorrhagic disease type 2 virus (RHDV2) in Sicily from rabbit (*Oryctolagus cuniculus*) and Italian hare (*Lepus corsicanus*). Research in Veterinary Science 2014;97(3):642–645 10.1016/j.rvsc.2014.10.008.

7. Hall RN, Peacock DE, Kovaliski J, Mahar JE, Mourant R et al. Detection of RHDV2 in European brown hares (*Lepus europaeus*) in Australia. Veterinary Record 2017;180(5): 121 10.1136/vr.104034.

8. Lankton JS, Knowles S, Keller S, Shearn-Bochsler VI, Ip HS. Pathology of *Lagovirus europaeus* GI.2/RHDV2/b (Rabbit hemorrhagic disease virus 2) in native North American lagomorphs. Journal of Wildlife Diseases 2021;57(3):694–700 10.7589/JWD-D-20-00207.

9. Le Gall-Recule G, Lemaitre E, Bertagnoli S, Hubert C, Top S et al. Large-scale lagovirus disease outbreaks in European brown hares (*Lepus europaeus*) in France caused by RHDV2 strains spatially shared with rabbits (*Oryctolagus cuniculus*). Veterinary Research 2017;48(1):70 10.1186/s13567-017-0473-y.

10. Neimanis AS, Ahola H, Larsson Pettersson U, Lopes AM, Abrantes J et al. Overcoming species barriers: an outbreak of *Lagovirus europaeus* GI.2/RHDV2 in an isolated population of mountain hares (*Lepus timidus*). BMC Veterinary Research 2018;14(1):367 10.1186/s12917-018-1694-7.

11. Puggioni G, Cavadini P, Maestrale C, Scivoli R, Botti G et al. The new French 2010 rabbit hemorrhagic disease virus causes an RHD-like disease in the Sardinian Cape hare (*Lepus capensis mediterraneus*). Veterinary Research 2013;44:96 10.1186/1297-9716-44-96.

12. Velarde R, Cavadini P, Neimanis A, Cabezon O, Chiari M et al. Spillover events of infection of Brown hares (*Lepus europaeus*) with rabbit haemorrhagic disease type 2 virus (RHDV2) caused sporadic cases of an European Brown Hare syndrome-like disease in Italy and Spain. Transboundary and Emerging Diseases 2017;64(6): 1750-1761 10.1111/tbed.12562.

13. Konig M, Thiel HJ, Meyers G. Detection of viral proteins after infection of cultured hepatocytes with rabbit hemorrhagic disease virus. Journal of Virology 1998;72(5):4492–4497 10.1128/JVI.72.5.4492-4497.1998.

14. Liu G, Zhang Y, Ni Z, Yun T, Sheng Z et al. Recovery of infectious rabbit hemorrhagic disease virus from rabbits after direct inoculation with *in vitro*-transcribed RNA. Journal of Virology 2006;80(13):6597–6602 10.1128/JVI.02078-05.

15. Liu G, Ni Z, Yun T, Yu B, Chen L et al. A DNA-launched reverse genetics system for rabbit hemorrhagic disease virus reveals that the VP2 protein is not essential for virus infectivity. Journal of General Virology 2008;89(12):3080–3085 10.1099/vir.0.2008/003525-0.

16. Wang B, Zhe M, Chen Z, Li C, Meng C et al. Construction and applications of rabbit hemorrhagic disease virus replicon. PLoS One 2013;8(5):e60316 10.1371/journal.pone.0060316.

17. Zhu J, Miao Q, Tan Y, Guo H, Liu T et al. Inclusion of an Arg-Gly-Asp receptor-recognition motif into the capsid protein of rabbit hemorrhagic disease virus enables culture of the virus *in vitro*. Journal of Biological Chemistry 2017;292(21):8605–8615 10.1074/jbc.M117.780924.

18. Ettayebi K, Crawford SE, Murakami K, Broughman JR, Karandikar U et al. Replication of human noroviruses in stem cell-derived human enteroids. Science 2016;353(6306):1387–1393 10.1126/science.aaf5211.

19. Cullen JM, Stalker MJ. Chapter 2 - Liver and Biliary System. In: Maxie MG (editor). Jubb, Kennedy & Palmer’s Pathology of Domestic Animals: Volume 2 (Sixth Edition): W.B. Saunders; 2016. pp. 258–352.e251 10.1016/B978-0-7020-5318-4.00008-5.

20. Vekemans K, Braet F. Structural and functional aspects of the liver and liver sinusoidal cells in relation to colon carcinoma metastasis. World Journal of Gastroenterology 2005;11(33):5095–5102 10.3748/wjg.v11.i33.5095.

21. Huch M, Gehart H, van Boxtel R, Hamer K, Blokzijl F et al. Long-term culture of genome-stable bipotent stem cells from adult human liver. Cell 2015;160(1-2):299–312 10.1016/j.cell.2014.11.050.

22. Sugiyama Y, Koike T, Shiojiri N. Immunohistochemical analyses of cell-cell interactions during hepatic organoid formation from fetal mouse liver cells cultured *in vitro*. Histochemistry and Cell Biology 2007;128(6):521–531 10.1007/s00418-007-0339-x.

23. Gabriel V, Zdyrski C, Sahoo DK, Dao K, Bourgois-Mochel A et al. Standardization and maintenance of 3D canine hepatic and intestinal organoid cultures for use in biomedical research. Journal of Visualized Experiments 2022(179) 10.3791/63515.

24. Lee JH, Lee DH, Park JK, Kim SK, Kwon CH et al. Potentiality of immobilized pig hepatocyte spheroids in bioartificial liver system. Transplantation Proceedings 2012;44(4): 1012-1014 10.1016/j.transproceed.2012.03.010.

25. Kruitwagen HS, Oosterhoff LA, Vernooij I, Schrall IM, van Wolferen ME et al. Long-term adult feline liver organoid cultures for disease modeling of hepatic steatosis. Stem Cell Reports 2017;8(4):822–830 10.1016/j.stemcr.2017.02.015.

26. Baron MG, Mintram KS, Owen SF, Hetheridge MJ, Moody AJ et al. Pharmaceutical metabolism in fish: Using a 3-D hepatic *in vitro m*odel to assess clearance. PLoS One 2017;12(1):e0168837 10.1371/journal.pone.0168837.

27. Baquerre C, Montillet G, Pain B. Liver organoids in domestic animals: an expected promise for metabolic studies. Veterinary Research 2021;52(1):47 10.1186/s13567-021-00916-y.

28. Broutier L, Andersson-Rolf A, Hindley CJ, Boj SF, Clevers H et al. Culture and establishment of self-renewing human and mouse adult liver and pancreas 3D organoids and their genetic manipulation. Nature Protocols 2016;11(9): 1724–1743 10.1038/nprot.2016.097.

29. Blutt SE, Crawford SE, Bomidi C, Zeng XL, Broughman JR et al. Use of human tissue stem cell-derived organoid cultures to model enterohepatic circulation. Am J Physiol Gastrointest Liver Physiol 2021;321(3):G270–G279 10.1152/ajpgi.00177.2021.

30. Huch M, Dorrell C, Boj SF, van Es JH, Li VS et al. *In vitro* expansion of single Lgr5+ liver stem cells induced by Wnt-driven regeneration. Nature 2013;494(7436):247–250 10.1038/nature11826.

31. Kardia E, Frese M, Smertina E, Strive T, Zeng XL et al. Culture and differentiation of rabbit intestinal organoids and organoid-derived cell monolayers. Scientific Reports 2021;11(1):5401 10.1038/s41598-021-84774-w.

32. Read AJ, Kirkland PD. Efficacy of a commercial vaccine against different strains of rabbit haemorrhagic disease virus. Australian Veterinary Journal 2017;95(7):223–226 10.1111/avj.12600.

33. Hall RN, Mahar JE, Read AJ, Mourant R, Piper M et al. A strain-specific multiplex RT-PCR for Australian rabbit haemorrhagic disease viruses uncovers a new recombinant virus variant in rabbits and hares. Transboundary and Emerging Diseases 2018;65(2):e444–e456 10.1111/tbed.12779.

34. Urakova N, Frese M, Hall RN, Liu J, Matthaei M et al. Expression and partial characterisation of rabbit haemorrhagic disease virus non-structural proteins. Virology 2015;484:69–79 10.1016/j.virol.2015.05.004.

35. Adams JM, Huppert KA, Castro EC, Lopez MF, Niknejad N et al. Sox9 Is a modifier of the liver disease severity in a mouse model of Alagille syndrome. Hepatology 2020;71(4): 1331-1349 10.1002/hep.30912.

36. Verstegen MMA, Roos FJM, Burka K, Gehart H, Jager M et al. Human extrahepatic and intrahepatic cholangiocyte organoids show region-specific differentiation potential and model cystic fibrosis-related bile duct disease. Scientific Reports 2020;10(1):21900 10.1038/s41598-020-79082-8.

37. Urakova N, Hall R, Strive T, Frese M. Restricted host specificity of rabbit hemorrhagic disease virus is supported by challenge experiments in immune-compromised mice (*Mus musculus*). Journal of Wildlife Diseases 2019;55(1):218–222 10.7589/2018-03-067.

38. Forootan A, Sjoback R, Bjorkman J, Sjogreen B, Linz L et al. Methods to determine limit of detection and limit of quantification in quantitative real-time PCR (qPCR). Biomolecular Detection and Quantification 2017;12:1–6 10.1016/j.bdq.2017.04.001.

39. Chinsangaram J, Hammami S, Osburn BI. Detection of bluetongue virus using a cDNA probe derived from genome segment 4 of bluetongue virus serotype 2. Journal of Veterinary Diagnostic Investigation 1992;4(1):8–12 10.1177/104063879200400103.

40. Ettayebi K, Tenge VR, Cortes-Penfield NW, Crawford SE, Neill FH et al. New insights and enhanced human norovirus cultivation in human intestinal enteroids. mSphere 2021;6(1) 10.1128/mSphere.01136-20.

41. Hosmillo M, Chaudhry Y, Nayak K, Sorgeloos F, Koo BK et al. Norovirus replication in human intestinal epithelial cells is restricted by the interferon-induced JAK/STAT signaling pathway and RNA polymerase II-mediated transcriptional responses. mBio 2020;11(2) 10.1128/mBio.00215-20.

42. Fiorotto R, Amenduni M, Mariotti V, Fabris L, Spirli C et al. Liver diseases in the dish: iPSC and organoids as a new approach to modeling liver diseases. Biochimica et Biophysica Acta Molecular Basis of Disease 2019;1865(5):920–928 10.1016/j.bbadis.2018.08.038.

43. Merchan T, Rocha G, Alda F, Silva E, Thompson G et al. Detection of rabbit haemorrhagic disease virus (RHDV) in nonspecific vertebrate hosts sympatric to the European wild rabbit (*Oryctolagus cuniculus*). Infection, Genetics and Evolution 2011;11(6): 1469–1474 10.1016/j.meegid.2011.05.001.

44. Rocha G, Alda F, Pages A, Merchan T. Experimental transmission of rabbit haemorrhagic disease virus (RHDV) from rabbit to wild mice (*Mus spretus* and *Apodemus sylvaticus*) under laboratory conditions. Infection, Genetics and Evolution 2017;47:94–98 10.1016/j.meegid.2016.11.016.

45. Calvete C, Mendoza M, Sarto MP, Bagues MPJ, Lujan L et al. Detection of rabbit hemorrhagic disease virus GI.2/RHDV2/b in the Mediterranean pine vole (*Microtus duodecimcostatus*) and white-toothed shrew (*Crocidura russula*). Journal of Wildlife Diseases 2019;55(2):467–472 10.7589/2018-05-124.

46. Abade Dos Santos FA, Pinto A, Burgoyne T, Dalton KP, Carvalho CL et al. Spillover events of rabbit haemorrhagic disease virus 2 (recombinant GI.4P-GI.2) from Lagomorpha to Eurasian badger. Transboundary and Emerging Diseases 2021 10.1111/tbed.14059.

47. Frolich K, Klima F, Dedek J. Antibodies against rabbit hemorrhagic disease virus in free-ranging red foxes from Germany. Journal of Wildlife Diseases 1998;34(3):436–442 10.7589/0090-3558-34.3.436.

48. Henning J, Meers J, Davies PR, Morris RS. Survival of rabbit haemorrhagic disease virus (RHDV) in the environment. Epidemiology and Infection 2005;133(4):719–730 10.1017/s0950268805003766.

49. Parkes JP, Heyward RP, Henning J, Motha MX. Antibody responses to rabbit haemorrhagic disease virus in predators, scavengers, and hares in New Zealand during epidemics in sympatric rabbit populations. New Zealand Veterinary Journal 2004;52(2):85–89 10.1080/00480169.2004.36410.

50. Gould AR, Kattenbelt JA, Lenghaus C, Morrissy C, Chamberlain T et al. The complete nucleotide sequence of rabbit haemorrhagic disease virus (Czech strain V351): use of the polymerase chain reaction to detect replication in Australian vertebrates and analysis of viral population sequence variation. Virus Research 1997;47(1):7–17 10.1016/s0168-1702(96)01399-8.

51. Zheng T, Lu G, Napier AM, Lockyer SJ. No virus replication in domestic cats fed with RHDV-infected rabbit livers. Veterinary Microbiology 2003;95(1-2):61–73 10.1016/s0378-1135(03)00135-4.

52. Sjostedt E, Zhong W, Fagerberg L, Karlsson M, Mitsios N et al. An atlas of the protein-coding genes in the human, pig, and mouse brain. Science 2020;367(6482) 10.1126/science.aay5947.

